# Sleep Deprivation Induces Acute Dissociation via Altered EEG Rhythms Expression and Connectivity

**DOI:** 10.1101/2022.03.21.485177

**Authors:** Danilo Menicucci, Valentina Cesari, Enrico Cipriani, Andrea Piarulli, Angelo Gemignani

## Abstract

The fragmented sleep, fragmented mind hypothesis has associated sleep disturbances and dissociative states in subjects with dissociative traits, as supported by neurophysiological theories of consciousness stating that altered states might result from an altered functional interaction among brain modules due to inefficient sleep processes.

Irrespective of dissociative traits, it is conceivable that a labile sleep-wake cycle might fuel dissociative states such as derealization, depersonalization, and dissociative amnesia.

To verify whether acute sleep loss can prompt dissociative states and to identify possible psychophysiological correlates, we evaluated dissociative experiences (by means of Phenomenology of Consciousness Inventory and Clinician Administered Dissociative State Scale) and resting state EEG features (band-wise spectral content and phase synchronization) after total sleep deprivation.

After deprivation, participants reported increased perception of altered state of consciousness and dissociative experiences, and a decreased perception of cognitive control. Analyzing the psychophysiological correlates of dissociative states following deprivation, we observed the following results: the higher the prefrontal theta spectral content, the higher the depersonalization state and the lower the self-awareness; the higher the intensity of the dissociative experiences, the higher the synchronization increase in alpha, beta, and gamma bands; the higher the decrease of higher-order functions, the higher the synchronization in the aforementioned bands.

Thus, acute sleep deprivation appears to fuel dissociative experiences by establishing a state of consciousness promoted by a higher large-scale synchronization at high frequencies.

## Introduction

Moment by moment, conscious content manifests itself as a unified scene generated from stimuli, memories, and their interplay; it is the continuous stream changing on a time scale of fractions of seconds by which William James described consciousness (James, 1892). As Gerald Edelman proposed in a series of seminal studies, the reentry between modality-specific cortical areas (posteriorly) and areas devoted to memory and executive functions (anteriorly) allows for the rise of conscious processes, thus sustaining the formation of a Dynamic Core; “a large cluster of neuronal groups that together constitute, on a time scale of hundreds of milliseconds, a unified neural process of high complexity” (Tononi and Edelman, 1998). Edelman hypothesized that sequences of integrated activity give rise to the unitary scenes that constitute the phenomenal experience (Edelman et al., 1992). The observation that the Dynamic Core (Edelman and Tononi, 2000) involves the integration of activity in widespread distributed cortical areas sustains the Global Workspace theory (Baars, 1997, 2002). The Global Workspace theory reconciles the limited capacity of momentary conscious content with the vast repertoire of the unconscious mind including long-term memory. A global broadcast of focal conscious contents can be viewed as a momentary “snapshot” of the Dynamic Core activity.

According to the Global workspace theory, the cerebral hemispheres hold a “global workspace”, which metaphorically can be depicted as the “headquarters” of the cortex in which information is received from, and projects to, many cortical modules (i.e., brain areas). Importantly, only information processed by the global workspace reaches consciousness. The global neuronal workspace theory (Dehaene & Naccache, 2001), posits that a large variety of brain perceptual and non-perceptual modules can be mobilized into the global workspace and their outputs integrated, favoring the emergence of a conscious experience. In turn, at the microscopic level, each area/module contains a complex anatomical circuitry that supports a diversity of activity patterns. The repertoire of possible contents of consciousness is thus characterized by an enormous combinatorial diversity: each workspace state is ‘highly differentiated’ and of ‘high complexity’ (as Tononi and Edelman, 1998, termed it). Along this line, Tononi’s integrated information theory (IIT) conceives consciousness as integrated information that increases in proportion to a system’s ability to integrate information and to the availability within the system of a vast repertoire of differentiated information. Higher values of complexity tend to reflect a higher degree of functional integration and specialization within the brain system, whereas elements within the system, which lead to a variety of less integrated patterns, typically result in decreased neural complexity (Tononi, 2004). Recently, the proponents of this theory developed a phenomenological frame of reference for conscious experience. This framework has been concisely systematized by the identification of six axioms that capture the subjective aspects of consciousness experience (hard problem, Chalmers, 1996).

When these aforementioned axioms are violated, consciousness can be subjected to various degrees of deviation from ordinary functioning, the so-called “altered states of consciousness” (Tart, 1975; Tononi et al., 2016).

The occurrence of an altered state might be partially explained either by a reduction of large-scale integration or by the reduction of the available repertoire of information (differentiation) within the brain. The level of independence among these complex, interlinked, and simultaneously active neural states, is reflected in neural complexity that takes into account the number of variables in determining the state of the system (Boly et al., 2009). Altered states of consciousness have been found in association with a disruption of the information integration in the brain (Alkire et al., 2008; Lee et al., 2009; Tononi, 2010; Tononi and Koch, 2015); the flow and the phenomenological characteristics of consciousness incur notable changes when an altered state arises (Edelman et al., 2011). Furthermore, temporal coordination is an obligatory condition for brain areas to integrate and transmit information; the recovery pattern of the brain network synchronization may reflect the recovery pattern of consciousness (Kim et al., 2018).

It has been speculated that certain functional disorders of consciousness, most notably dissociative disorders, might be characterized by the formation of multiple dynamic cores operating with high levels of independence, without a precise and common spatio-temporal frame of reference (Lutzenberger et al., 1992). A similar theoretic contribution derives from the global workspace theory; in it multiple subunits of brain elaboration compete for the access to the main serial processor. In normal conditions, this competition is regulated; in altered states of consciousness this regulation may be disrupted, thus leading to an opposition between subunits and subsequently to altered informational elaboration (Baars, 1993).

In line with these hypotheses, dissociation is defined as “characterized by a disruption of, and/or discontinuity in the normal integration of consciousness” (American Psychiatric Association, 2013, p.291), thus resulting in derealization, depersonalization, absorption and dissociative amnesia. Dissociative experiences represent the core symptoms of dissociative disorders (American Psychiatric Association, 2013). They occur in a wide range of psychopathological and somatic diseases, but also in non-clinical populations (Lyssenko et al., 2018). Dissociation has several hallmarks emerging from prototypical EEG-derived features. For example, EEG theta and delta power increases, together with alpha power decreases, are found to be positively correlated to dissociation during resting state (Giesbrecht et al., 2006; Mesulam, 1981; Sierra and Berrios, 1998). From a connectivity standpoint, the fragmented state typical of dissociative experience is characterized by higher levels of segregation and lower levels of integration of information between brain areas. Indeed Soffer-Dudek and colleagues (2018), observed that dissociative experiences were characterized by decrease of short (central-parietal) and long range (fronto-occipital) EEG coherence within delta and theta band.

However, trait dissociation does not represent a *conditio sine qua non* for the emergence of these experiences. In fact, state dissociation, which refers to a temporary dissociative state, could be also triggered by intense emotions, pharmacological inductions, or even by a labile sleep–wake cycle. This can occur even in the absence of a stable background trait. Along this line, some studies reported that alterations in the sleep-wake cycle may decrease memory and attentional control, promoting absentmindedness and a propensity to produce memory errors, two well-established correlates of dissociative states (Giesbrecht & Merckelback, 2004, Giesbrecht et al., 2010, Merckelbach et al., 2007).

Given the association between sleep disturbances and dissociative states, aptly named the “fragmented sleep, fragmented mind” hypothesis (Van der Kloet et al., 2012), the experimental manipulation of sleep through sleep deprivation is a reliable paradigm to fuel dissociative experiences in both clinical population and healthy participants.

If dissociative symptoms can be fueled by a labile sleep–wake cycle, then sleep loss would be expected to intensify dissociative symptoms (Morgan et al. 2001; Giesbrecht, et al., 2007). However, two importants issues arise from previous studies: the first is that previous works investigated sleep-dissociation by studying trait characteristic and sleep habits without directly manipulating the sleep variable, thus the study of sleep deprivation-induced acute dissociation (briefly, deprivation-induced acute dissociation) remained precluded; the second is that resulting outcomes pertained the effect of dissociative state on task-related performances (i.e., Schabinger et al., 2018) and cognitive domains, thus, due to its spontaneous and subjective nature, the phenomenological experience of dissociative states still remains uninvestigated. Indeed, the direct manipulation of sleep could offer the possibility to study deprivation-induced acute dissociation.

In this study we aim to verify whether sleep deprivation effectively increases dissociative experiences, namely derealization, depersonalization, and dissociative amnesia, and we identify and trace their electrophysiological correlates. We will then identify associations between intrasubjective variability from baseline to sleep deprivation of electrophysiological indices and measures of dissociation.

## Results

The dissociative-like experiences associated with sleep loss were neuro-phenomenologically characterized in 18 healthy subjects (9 males, 7 females, age 28 ± 7 years). We performed a *within-subject* study (experimental protocol in Figure 6) using sleep deprivation to induce changes in subjective conscious experiences. Volunteers underwent two randomized and balanced (across participants) experimental sessions at a one-week interval one from the other:

- baseline session: each participant was tested at 8 pm after a normal night of sleep
- deprivation session: each participant was tested at 8 pm after 36 hours of sleep deprivation.

Within each session, EEG signals were recorded during 10 minutes of eyes-closed resting state. Immediately after, a psychometric assessment was conducted with the aim of identifying dissociative-like experiences (such as perceiving the surrounding environment as “unreal”). The psychometric evaluation assessing perceived state of consciousness was carried out using the following instruments: the Phenomenology of Consciousness Inventory (PCI, Pekala, 1991) and the Clinician Administered Dissociative State Scale (CADSS, Bremner et al., 1998). Descriptive statistics of PCI and CADSS scores are reported in the *supplementary materials*.

From the EEG, we estimated features belonging to two classes: power spectral densities (PSD), and phase synchronizations (dWPLI) in five bands of interest. For each frequency band, PSD was estimated for each EEG channel, whereas phase synchronization was estimated between each couple of EEG channels.

For each psychometric measure, between-conditions differences (post-deprivation versus baseline) were computed (CADSS total score and its three subdimensions scores as well as PCI major dimensions) using the paired Wilcoxon Signed rank test (significance threshold was set at p ≤ 0.05). Furthermore, the Spearman’s rank correlation test was used to estimate associations between changes from baseline to post deprivation of significant psychometric measures and those of EEG features (PSD, dwPLI). Correlation significance was then evaluated using a single threshold permutation test for the maximum r-statistic with 10000 randomizations (Statistical nonParametric Mapping, SnPm, Nichols and Holmes, 2002).

All statistical analyses were computed using tailored codes written in Matlab (MathWorks, Natick, MA, USA) and Statistical Package for the Social Sciences version 25 (SPSS statistics, IBM SPSS, IBM Corporation).

For further details of experimental protocol, setup, and analysis, see Material and Methods sections.

### Dissociative experiences are strongly increased after sleep deprivation

Sleep deprivation condition was characterized by a significantly higher state dissociation (total CADSS score, p < 0.001) as compared to the control condition. When considering CADSS sub-dimensions, volunteers reported a higher level of depersonalization and derealization (p < 0.001 for both, see Table 1). The magnitude of effect sizes ranged from medium to large (Cohen et al., 1988).

**Table 1.** Clinician Administered Dissociative Symptom Scale (CADSS)

| **Name** | **z score** | **p value** | **Effect Size** |
| --- | --- | --- | --- |
| **Total Score** | 3.13 | **0.001\*\*\*** | **-0,54** |
| *Depersonalization* | 2.70 | **0.001\*\*\*** | **-0,46** |
| *Derealization* | 2.98 | **0.001\*\*\*** | **-0,51** |
| *Dissociative Amnesia* | 1.76 | 0.109 | -0,30 |
| *Wilcoxon signed rank test. \* $p < 0.05$, \*\* $p < 0.01$; \*\*\* $p < 0.001$* |  |  |  |

### Phenomenal facets of conscious experience are altered after sleep deprivation

Sleep deprivation was accompanied by significant changes in the subjective perception of many aspects of the resting state conscious experience when compared to the control session (Table 2). Volunteers reported a higher perception of having lived an altered experience (p < 0.05), accompanied by a lower level of self-awareness (p < 0.001), rationality, volition and memory (p < 0.01 for the formers and p < 0.001 for the latter), and a reduction of internal dialogue (p < 0.05). The magnitude of effect size ranged from medium to large (Cohen et al., 1988).

**Table 2.**
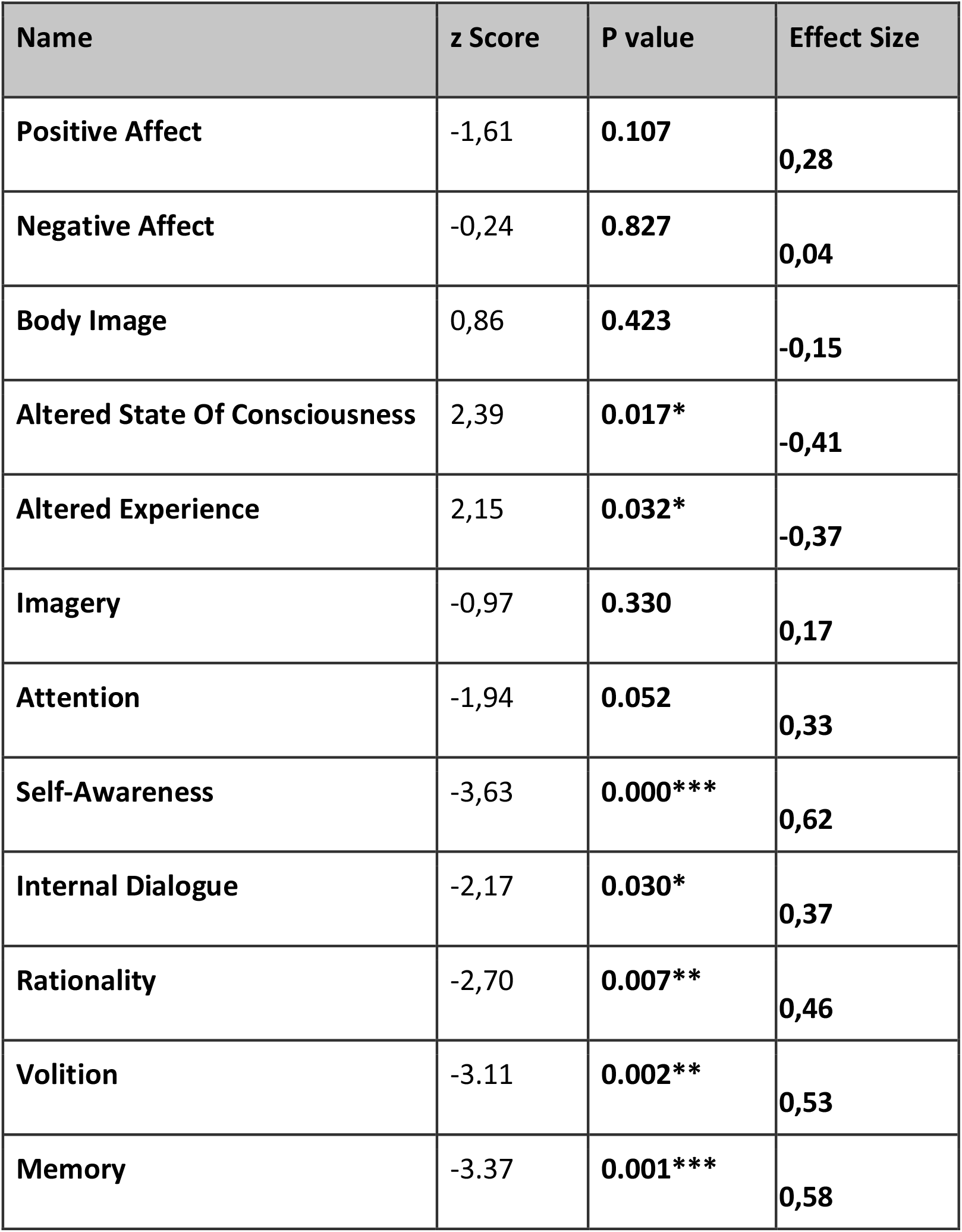
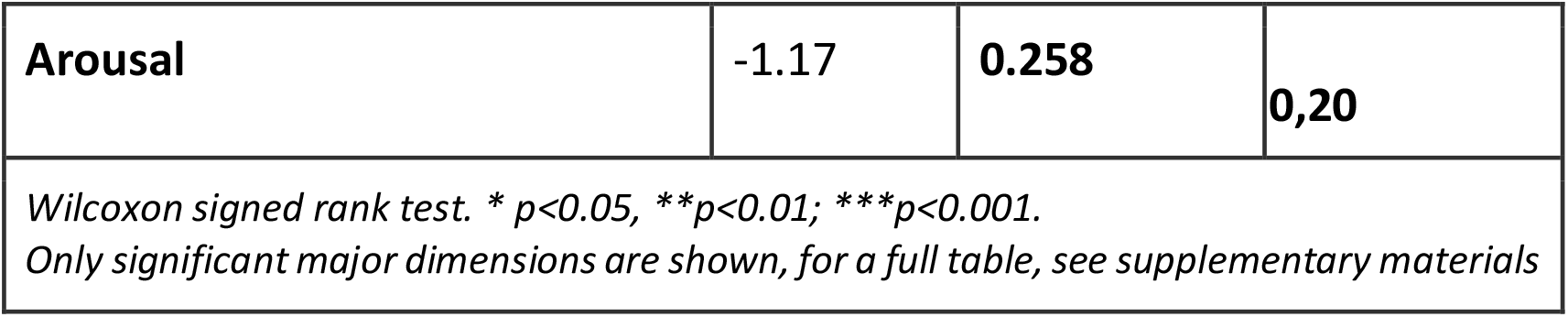
Phenomenology of Consciousness Inventory (PCI)

### The enhancement of frontal theta power is associated with a reduction of self awareness, after sleep deprivation

Bandwise associations between changes in EEG PSD and each psychometric measures (i.e., CADSS and PCI dimensions), were estimated for each EEG electrode; consequently, for each band and psychometric measure, correlation analysis yielded a map highlighting scalp areas associated with psychometric scores.

When considering theta band, we observed that between-condition variations in self awareness levels were negatively correlated with PSD changes over the whole scalp. This association was significant when considering prefrontal electrodes (p < 0.05 after SnPM correction): namely a reduction of self-awareness was significantly associated with an enhancement of theta PSD (Figure 1, Panel B).

**Figure 1:**
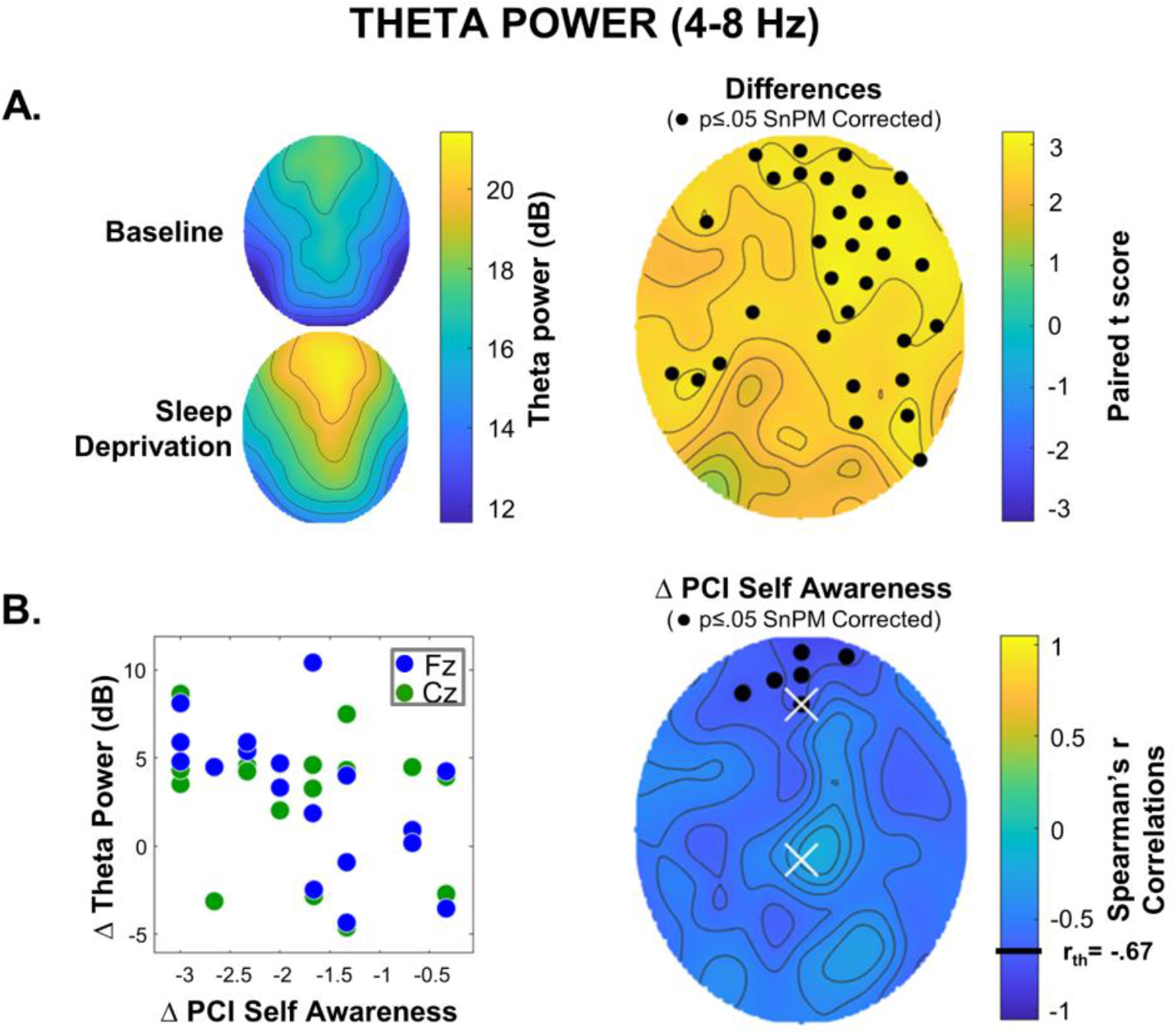
Theta power; inter-condition changes and correlation with changes in PCI Self Awareness. **Panel A:** Group averaged scalp maps of theta power in the baseline and sleep deprivation conditions, and their difference (electrodes for which a significant difference was detected after SnPM correction are marked with a black dot). After sleep deprivation theta power was significantly increased in electrodes spanning over the right frontal lobe and invading parietal areas. **Panel B**: On the right, scalp map of correlations between inter condition changes (Δ) of theta power and of PCI Self Awareness (electrodes for which a significant correlation was detected after SnPM correction are marked with black dots). On the left, scatterplot for the exemplary electrodes marked with a white X on the scalp map (right): each dot refers to a subject. Between-conditions changes in self awareness and theta power in the frontopolar area are negatively correlated.

Between-condition electrode-wise comparisons, showed an increase of theta PSD over the whole scalp after sleep deprivation as compared to control condition. The increase was significant (p<0.05 after SnPM correction) in bilateral prefrontal areas and in right fronto-central and posterior areas (Figure 1, Panel A).

### The enhancement of frontal gamma power is associated with an increase of depersonalization, after sleep deprivation

We next found a significant positive association (p < 0.05 after SnPM correction), between increases (sleep deprivation - control session) of gamma PSD in right fronto-lateral electrodes and concurrent increases in depersonalization. Indeed, subjects reporting a higher depersonalization after sleep deprivation were characterized at the EEG level by higher gamma band PSD in the right fronto-lateral scalp region (Figure 2, panel B).

**Figure 2:**
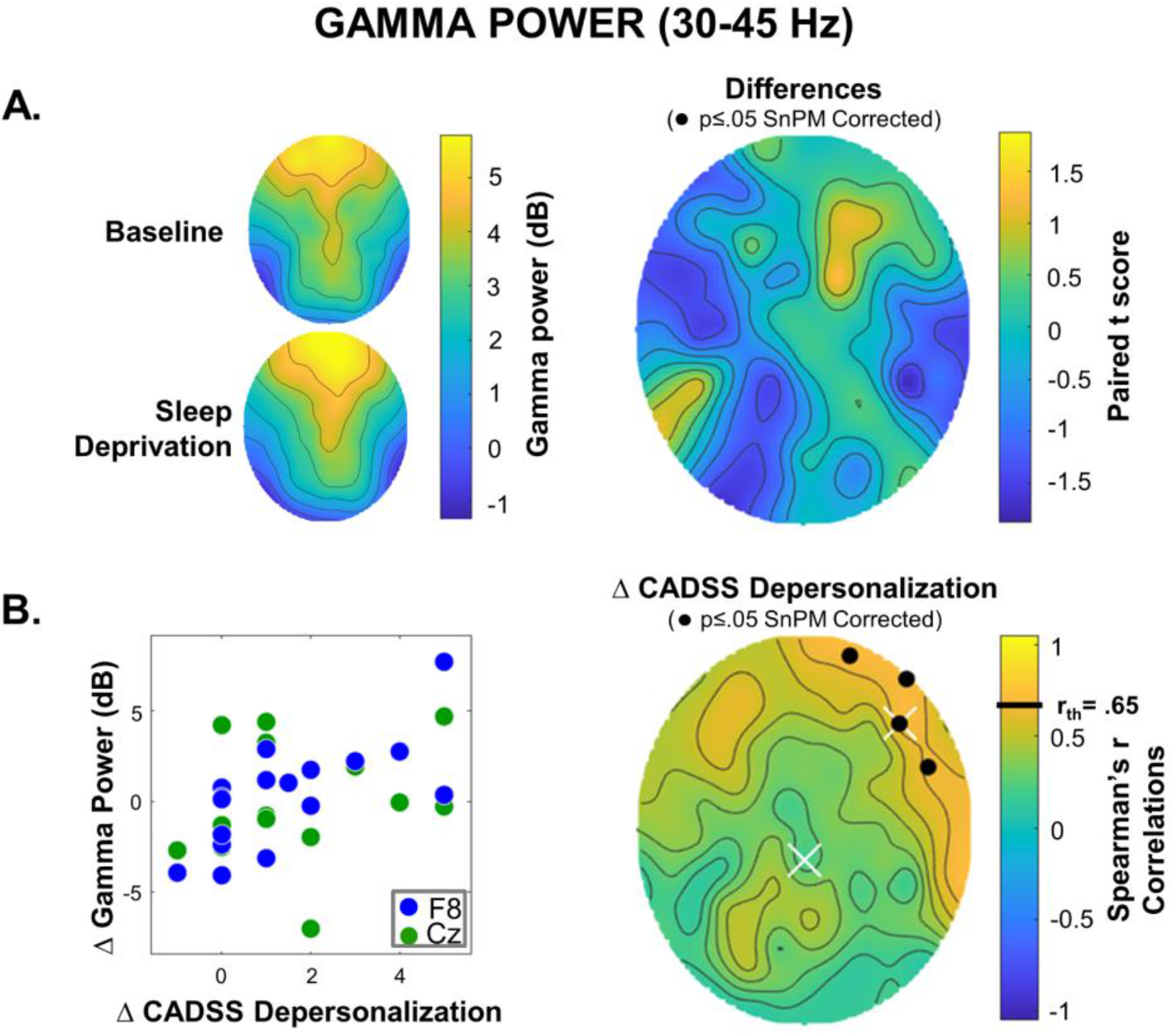
Gamma power; inter-condition changes and correlation with changes in CADSS Depersonalization. **Panel A:** Group averaged scalp maps of gamma power in the baseline and sleep deprivation conditions, and their difference. no significant differences were detected after SnPM correction. **Panel B**: On the right, scalp map of correlations between inter condition changes (Δ) of theta power and of Depersonalization (electrodes for which a significant correlation was detected after SnPM correction are marked with black dots). On the left, scatterplot for the exemplary electrodes marked with a white X on the scalp map (right): each dot refers to a subject. Between-conditions changes in depersonalization and gamma power in the right frontal cortex are correlated.

At variance with theta band, no significant between-condition difference (sleep deprivation versus baseline), was found for gamma band PSD (Figure 2, panel A).

### EEG phase synchronization highlights distinctive brain networks associated with dissociation and phenomenology of consciousness in sleep deprivation

Band-wise correlations between changes in EEG phase synchronization and variations in psychometric scores (CADSS and PCI) were calculated for each EEG electrode pair. Results of the correlation analyses are thus shown via scalp connectivity maps, where connectivities (between couples of electrodes) showing significant correlations with psychometric scores are represented as segments connecting the related electrodes.

#### Alpha Band

In the alpha band, total CADSS score changes from baseline to sleep deprivation correlated positively with connectivity variations between contralateral parietal areas, and between right centro-frontal and midline regions (Figure 3, panel A). Changes in the Derealization subscale were positively associated with connectivity changes in connectivity between contralateral parieto-occipital areas, right frontal and midline regions, and between left occipital area and the frontal pole (Figure 3, panel B). When considering PCI dimensions, we found significant positive associations between changes in Volitional Control and connectivity between left fronto-temporal and midline frontal areas (Figure 3, panel C), and between Altered Experience variations and connectivity changes between left occipital and right parietal regions (Figure 3, panel D). On the other hand, connectivity changes between right temporal and right frontal regions correlated negatively with changes in the Altered State of Awareness Dimension (Figure 3, panel E).

**Figure 3:**
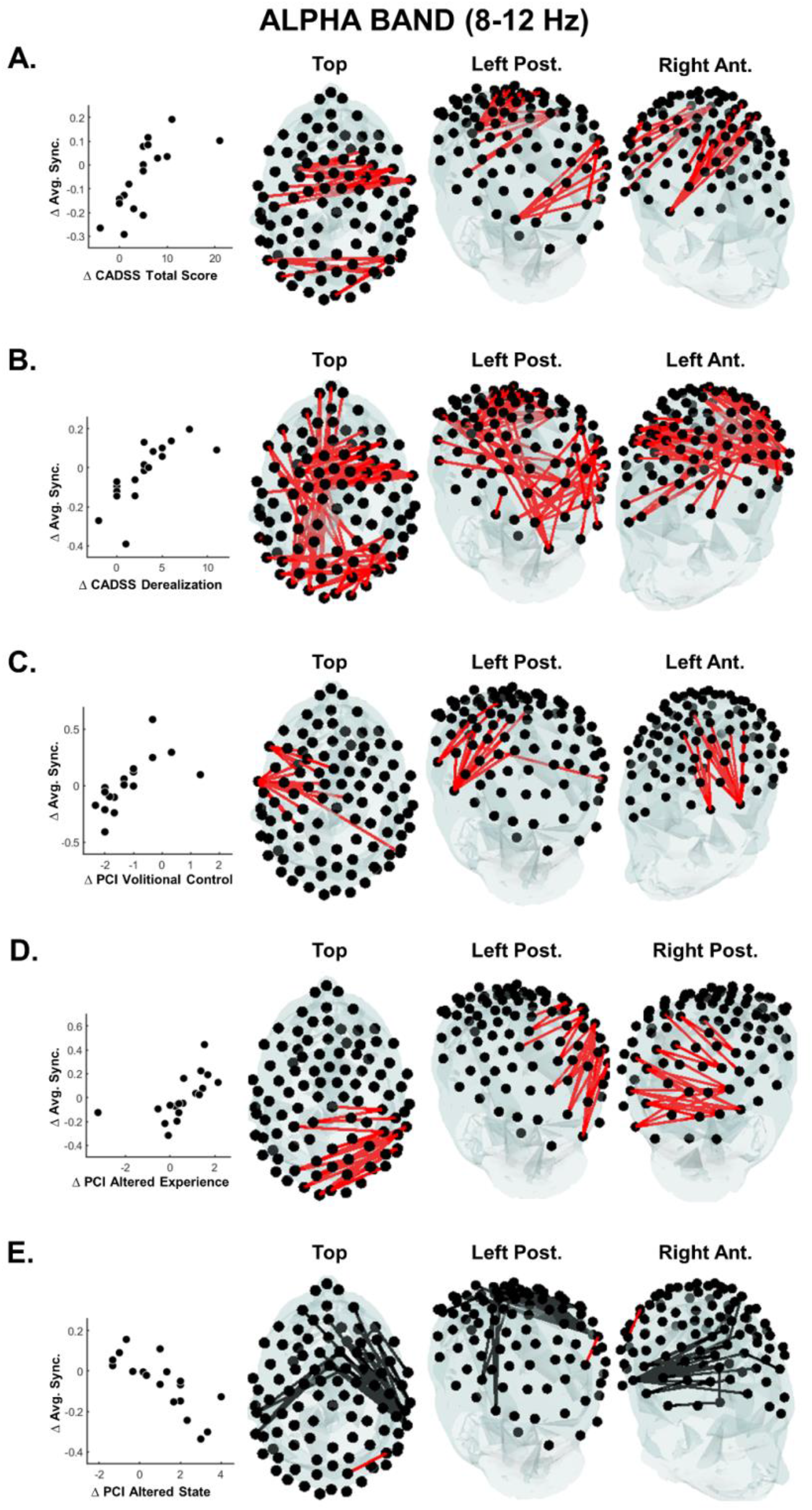
Correlations between EEG phase synchonization (DwPLI) in the alpha band and psychometric measures. Each panel refers to correlations with a psychometric measure. The correlations were calculated between the inter-condition changes of this psychometric measure and the synchronization changes measured for all possible electrode pairs (4851 pairs). The headplots show from multiple perspectives the set of electrode pairs whose synchronization change correlates significantly with the change in the psychometric measure (Spearman correlation, p<0.05 SnPM corrected). Red segments indicate a positive correlation, black ones a negative correlation. The scatterplot on the left highlights the average relationship between the set of significant electrode pairs and the psychometric measure. Each dot in the scatter plot represents a single subject and the y-axis value has been derived as the mean synchronization calculated over the electrode pairs with statistically significant correlation, as shown in the related headplots (Δ Avg. Sync). **Panel A:** Correlations between CADSS total score and DwPLI highlights bilateral connections between parietal and central-temporal cortices. **Panel B:** Correlations between CADSS Derealization score and DwPLI show bilateral connections between parietal and central-temporal areas, and between left occipital and the frontal pole. **Panel C:** Correlations between PCI Volitional control and DwPLI display connections between left fronto-temporal and midline frontal areas. **Panel D:** Correlations between PCI Altered Experience and DwPLI highlight connections between the right parietal cortex and midline posterior areas. **Panel E:** Correlations between PCI Altered State and DwPLI highlight connections between right temporal and right frontal areas.

No significant difference was found in alpha connectivity when comparing sleep deprivation to baseline.

#### Beta band

We observed a positive correlation between changes in total CADSS score and connectivity within occipital and temporal regions (Figure 4, panel A), and also between h changes in the Derealization subscale and connectivity among right occipito-parietal and left parietal regions (Figure 4, panel B). Regarding PCI scores,connectivity changes were negatively correlated with changes in the following dimensions:

**Figure 4:**
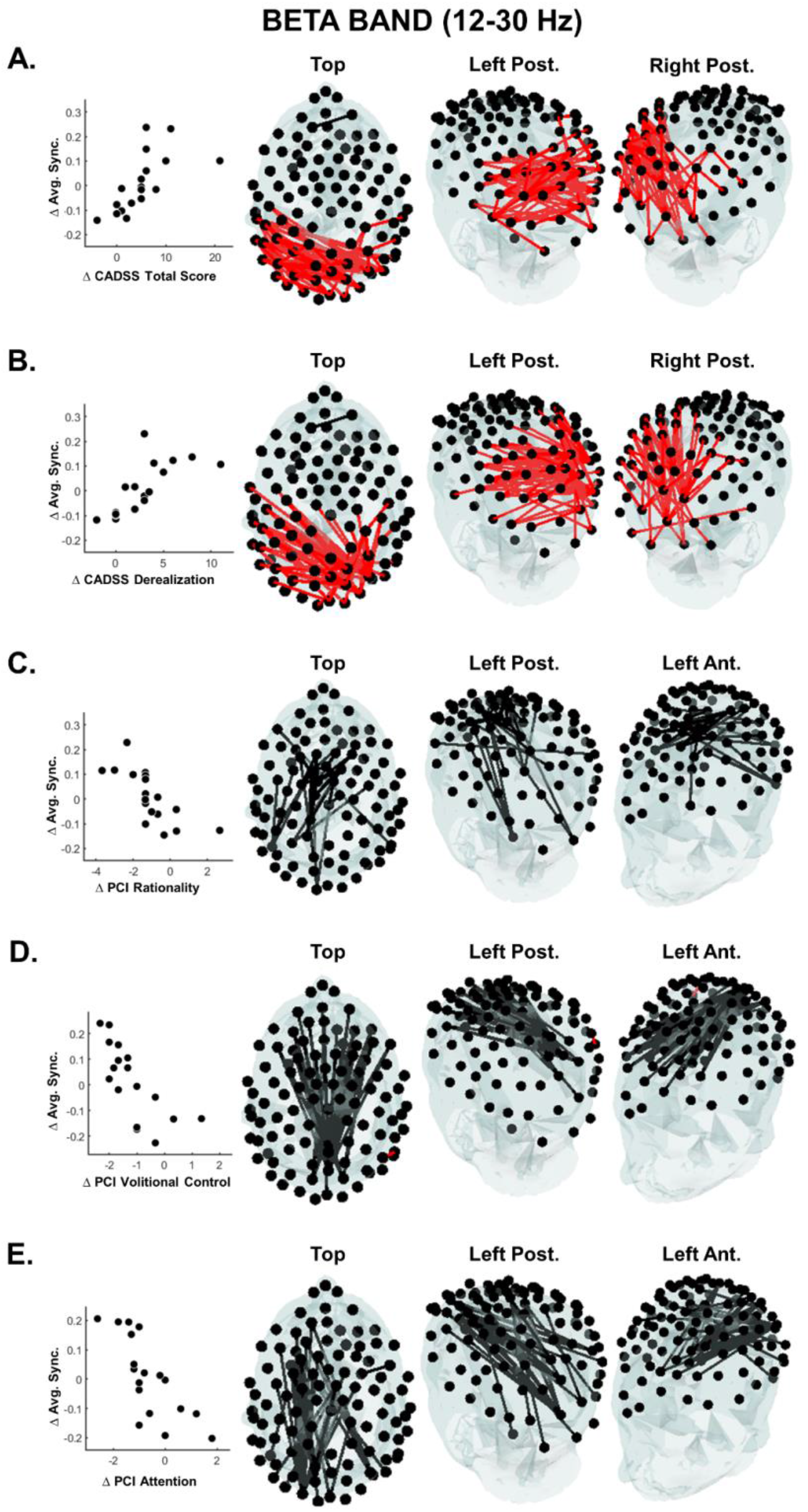
Correlations between EEG phase synchonization (DwPLI) in the beta band and psychometric measures. The correlations were calculated between the inter-condition changes of this psychometric measure and the synchronization changes measured for all possible electrode pairs (4851 pairs). The headplots show from multiple perspectives the set of electrode pairs whose synchronization change correlates significantly with the change in the psychometric measure (Spearman correlation, p<0.05 SnPM corrected). Red segments indicate a positive correlation, black ones a negative correlation. The scatterplot on the left highlights the average relationship between the set of significant electrode pairs and the psychometric measure. Each dot in the scatter plot represents a single subject and the y-axis value has been derived as the mean synchronization calculated over the electrode pairs with statistically significant correlation, as shown in the related headplots (Δ Avg. Sync). **Panel A:** Correlations between Total CADSS score and DwPLI highlight connections between bilateral occipital and temporal areas. **Panel B:** Correlations between CADSS Derealization score and DwPLI show connections between right occipito-parietal and left parietal areas. **Panel C:** Correlations between PCI Rationality and DwPLI display connections between left frontal and left parietal areas. **Panel D:** Correlations between PCI Volitional control and DwPLI show connections between a midline posterior parietal area and frontal cortices. **Panel E:** Correlations between PCI Attention and DwPLI highlight a bundle of connections between left frontal and left parietal cortex.

- Attention, connectivity between frontal and parieto-occipital regions with a left hemispheric prevalence (Figure 4, panel E);
- Volitional control, connectivity between midline parietal areas to widespread portions of the frontal regions (Figure 4, panel D);
- Rationality, connectivity within midline fronto-central areas cortex (Figure 4, panel C). In line with results regarding alpha band, no significant between-condition difference was found also for connectivity within beta band.

#### Gamma Band

In the gamma band, changes in the total CADSS score were positively correlated with variations of connectivity strength between right occipito-parietal areas and left temporal ones with (Figure 5, panel A). An analogous relationship was found also when considering changes in the Depersonalization subscale (Figure 5, panel B). Regarding PCI scores, we observed a negative correlation between changes in left temporal area to bilateral frontal regions connectivity and changes in Internal Dialogue (Figure 5, panel C). No significant between-condition difference was found between post-deprivation and baseline connectivity.

**Figure 5:**
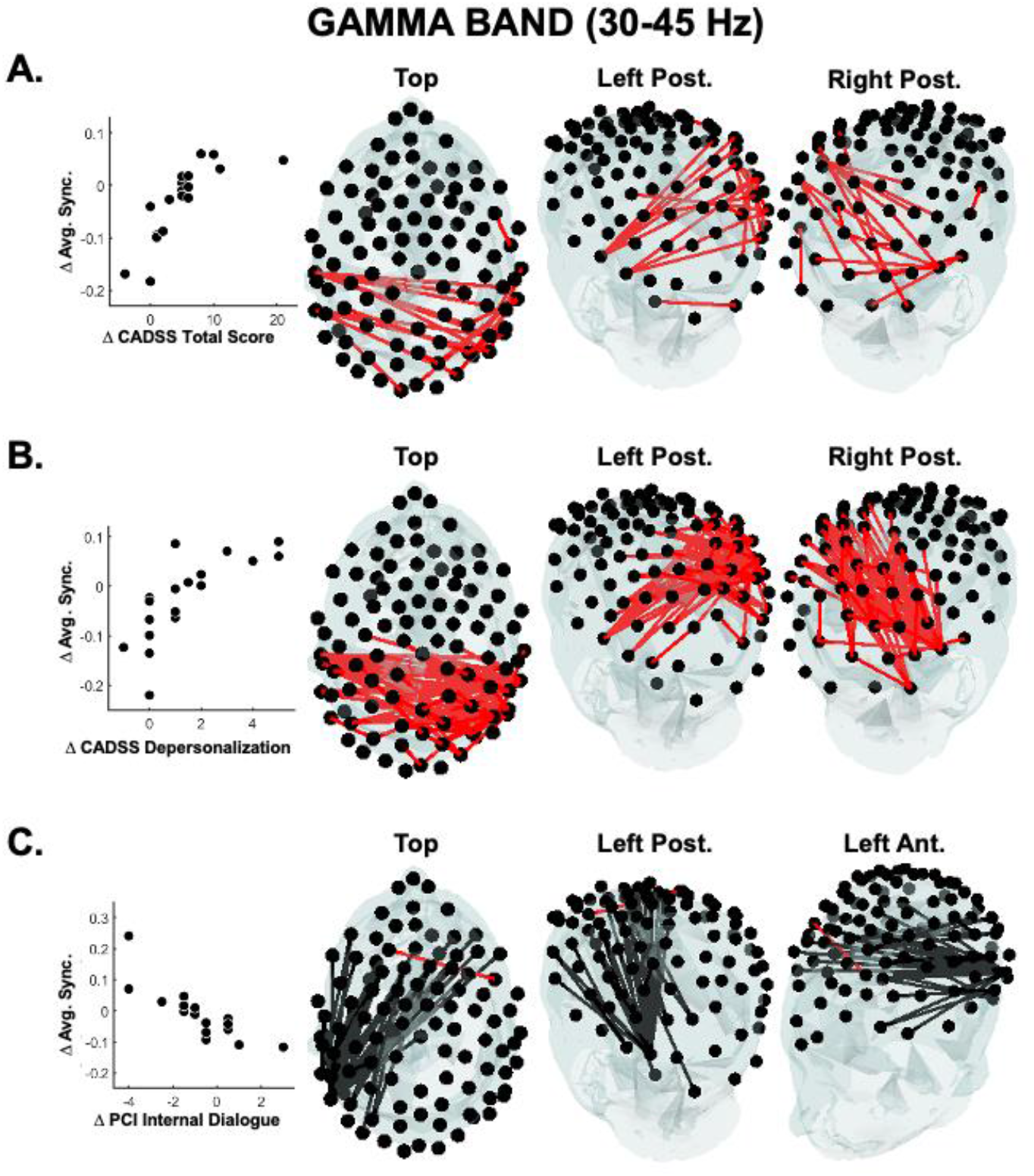
Correlations between EEG phase synchonization (DwPLI) in the gamma band and psychometric measures. Each panel refers to correlations with a psychometric measure. The correlations were calculated between the inter-condition changes of this psychometric measure and the synchronization changes measured for all possible electrode pairs (4851 pairs). The headplots show from multiple perspectives the set of electrode pairs whose synchronization change correlates significantly with the change in the psychometric measure (Spearman correlation, p<0.05 SnPM corrected). Red segments indicate a positive correlation, black ones a negative correlation. The scatterplot on the left highlights the average relationship between the set of significant electrode pairs and the psychometric measure. Each dot in the scatter plot represents a single subject and the y-axis value has been derived as the mean synchronization calculated over the electrode pairs with statistically significant correlation, as shown in the related headplots (Δ Avg. Sync). **Panel A:** Correlation between Total CADSS score and DwPLI highlights connections between bilateral temporal and parietal areas. **Panel B:** Correlation between CADSS Depersonalization score and DwPLI highlight connections between bilateral parietal and temporal areas. **Panel C:** Correlations between PCI Internal Dialogue and DwPLI display connections between left temporal and bilateral frontal areas.

Figures 4, 5, and 6 show the frequency bands in which we found significant results (significance thresholds corrected for multiple tests – 4851 electrode pairs – using the SnPM approach).

**Figure 6:**
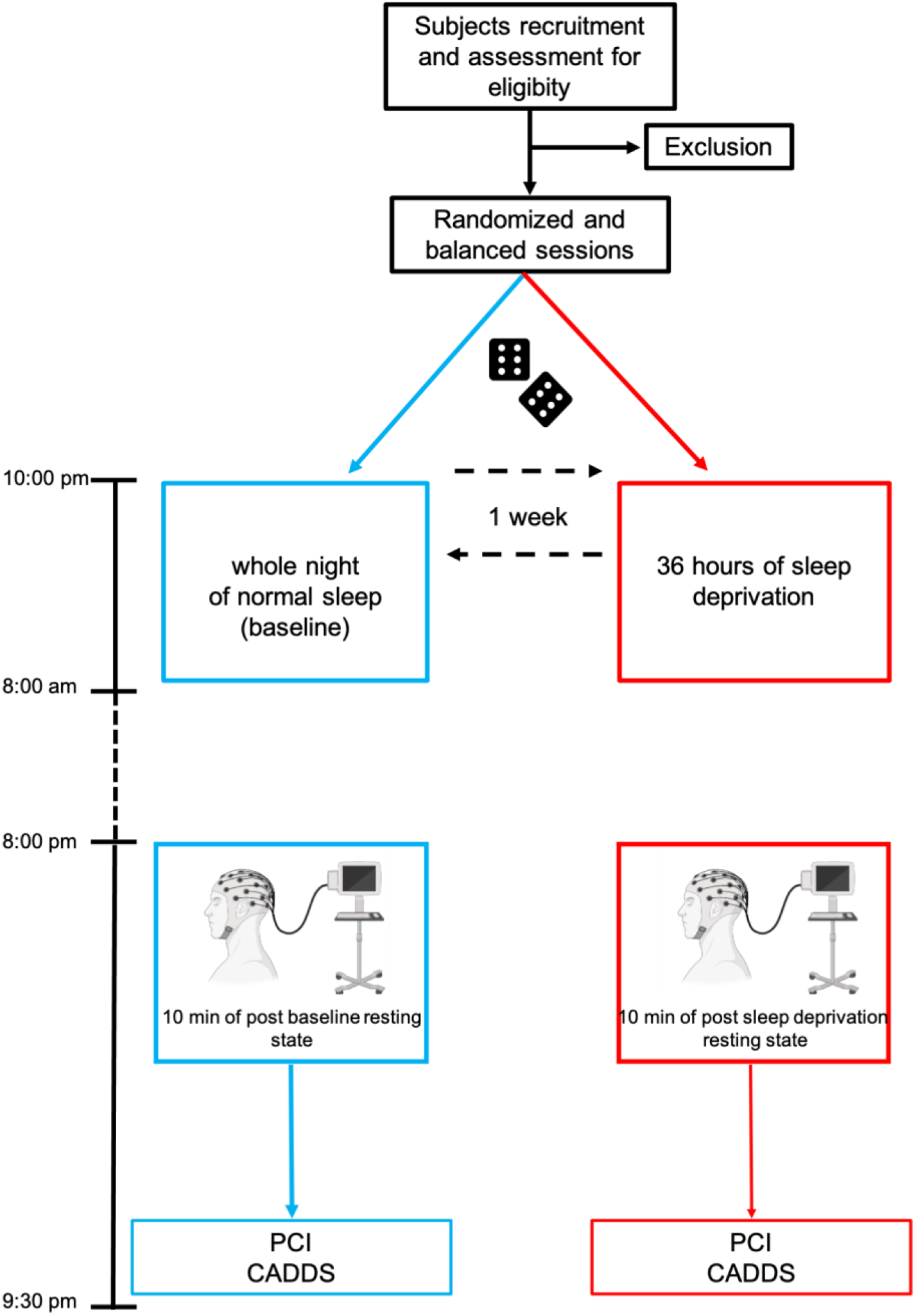
Flow chart of the experimental protocol. Timeline for both conditions (blue, baseline; red, post sleep deprivation).

## Discussion

The study investigated alterations in conscious experiences ascribable to a dissociative state emerging after sleep deprivation, and sought to identify the electrophysiological correlates of these alterations. After deprivation, participants reported increased perception of altered state of consciousness and dissociative experiences, and a decreased perception of cognitive control. Parallelly, participants displayed significant association between dissociative experiences and functional connectivity and band spectral content in high frequency.

Analyzing the psychophysiological correlates of dissociative states following deprivation, we observed the following results: the higher the prefrontal theta spectral content, the higher the depersonalization state and the lower the self-awareness; the higher the intensity of the dissociative experiences, the higher the synchronization increase in alpha, beta, and gamma bands; the higher the decrease of higher-order functions, the higher the synchronization in the aforementioned bands.

### Acute sleep manipulation and state characterization allow the conceptualization of a “deprivation-induced acute dissociation hypothesis”

We observed an increased perception of having experienced an altered state of consciousness during the resting state of the sleep deprivation session as compared to that of the baseline session. Subjects reported higher “pure” dissociative features (i.e., depersonalization and derealization). Moreover, taking in an extended conceptualization of “state dissociation”, the volunteers reported a a reduction in perceived self awareness, rationality of thoughts, and volitional control, paralleled with an increase in the subjective perception of being in an altered state of consciousness corresponding to commonly reported phenomenological features of dissociation (American Psychiatric Association, 2013). These findings are consistent with previous studies investigating the existing positive correlation between trait-dissociation and labile sleep cycle, corroborating the “*fragmented-sleep, fragmented mind*” hypothesis (van Der Kloet et al., 2012). In line with this well established association, we hypothesize that one night of sleep deprivation might prompt a state-dissociative experience. Thus, in an extended framework, this speculation could integrate the previous model, seminally elucidated by van Der Kloet and colleagues (2012).

### A “dissociative rebound system” might counteract sleep deprivation effects

We found a significant increase of theta band power after sleep deprivation as compared to baseline at prefrontal and right fronto-central sites. This result suggests that the enhancement of slow frequency bands PSD might be associated with a sleep deprivation main effect, in line with previous studies showing a predominance of delta and theta bands in frontal sites (Cajochen et al., 1999; Finelli et al., al., 2000; Strijkstra et al., 2003) as a marker of EEG-sleep propensity during wakefulness.

Theta band power increase was associated with a decrease of self-awareness from baseline to post sleep deprivation, whereas depersonalization increase was positively associated with an increase of gamma band power spectral density. Our findings about theta band negative association with self-awareness (a common feature of dissociative-like experiences), confirm and extend results from previous studies highlighting an association between theta and delta power synchronization and dissociation during resting states (Giesbrecht et al., 2006; Mesulam, 1981; Hollander et al., 1992; Sierra and Berrios, 1998). In an extended framework, negative associations between slow frequencies and awareness are found in a wide variety of conditions characterized by deviations from ordinary states of consciousness for instance, increased theta after ultraslow mechanical stimulation of nasal epithelium (Piarulli et al., 2018), or after hypnotic induction after relaxation techniques (Holroyd, 2003). Despite the heterogeneity of the aforementioned conditions, the enhancement of slow frequencies appears to be an EEG hallmark of altered consciousness in healthy populations.

For what concerns the gamma/depersonalization positive correlation, previous studies pertaining to dissociative literature did not find specific alterations of high frequency EEG activity, while studies on sleep deprivation found an overall decrease in gamma power density (Liet al., 2008). A possible explanation for the gamma/depersonalization positive association could be the rise of a “rebound-alertness state” which would partially counteract the effect of sleep pressure, in which the frontal gamma might play a pivotal role: indeed, studies on vigilance and sleep in humans have provided converging evidence for the association between gamma activity and states of arousal or alertness (Makeig and Jung, 1996). Another study found that the local effects in frontal gamma power might represent a compensatory mechanism to counter processing deficits (i.e., hyperfrontality), as gamma activity has been found to reflect local neuronal spiking and frontal activity is strongly linked to executive functions (Callicott et al., 2003; Manoach, 2003). Thus, the supposed increased level of alertness, and “hyperfrontality” might also account for the emergence of depersonalization experiences after sleep deprivation. In fact, depersonalization has been prototypically defined as a “thinking without feeling” condition (Phillips, 2001), and it has been hypothesized that depersonalization/derealization experiences might rise from a lack of integration between limbic, affective information, and cortical cognitive reasoning, in what has been dubbed as the “*Corticolimbic disconnection model*” of dissociation (Sierra and Berrios, 1998), on which we will return when discussing functional connectivity findings.

Another essential element of our results is the right hemisphere lateralization of power density changes (Figure 1 and 3). It has been reported that right regions, and in particular the right frontal ones, play a pivotal role in self-related functions (Feinberg, 2013). Our findings seem thus to suggest that the right frontal hemisphere susceptibility to sleep deprivation could entail alterations in self-awareness. On this basis we hypothesize that the deprivation-induced acute dissociation might be mainly explained by a subjective altered state of consciousness in which hyperfrontality, driven by frontal gamma, could represent a mechanism aiming at counteracting the emergence of drowsiness marked instead by theta power increases.

### Altered state experience associated with connectivity via phase synchronization of high rhythms

In our study we found multiple correlations between differences between the conditions in high frequency EEG bands (alpha, beta and gamma) phase synchronization. Phase synchronization appears to have a positive correlation with dissociation measures in all three aforementioned bands (Figure 3, panel A and B; Figure 4, panel A and B; Figure 5 panel A and B), while showing a negative correlation with psychometric measures relating to volitional control and attention (iFigure 3, panel C, D, and E; Figure 4, panel C, D, and E; Figure 5, panel C).

Different models can help describe these results within a unitary framework. These findings could be ascribed to the tendency of the brain to seek an optimal condition by which top-down frontal executive functions effectively regulate attention *via* inhibitory processes (Diamonds, 2013; Manuel et al. 2013; Chavan et al. 2015).

Results are also compatible with Tononi’s information integration theory (2008), which implies that the capacity to integrate information is reduced when neural activity becomes too synchronous; this could be ascribed to the violation of the axiom of integration postulating that consciousness is irreducible to non-interdependent components)

The present results could also be interpreted in the framework of the “*Deafferentation model”* (Newberg and D’Aquili, 2000), which might explain how altered connectivity could elicit altered states of consciousness such as dissociation. In this model, dissociation arises from the loss of input to higher-level associative brain regions. When these areas become cut-off from the rest of the brain due to altered connectivity, a loss of integration occurs; thus leading to a “*disruption of the normally integrated functions of consciousness, memory, identity, or perception of the environment*” (American Psychiatric Association, 2013) that is to the phenomenal experience of dissociation. As we will discuss later, our findings seem to highlight an alteration in connectivity between higher-order associative brain regions and the rest of the brain.

Another avenue of interpretation for these results might be derived from the theoretical framework dubbed the “information via desynchronization hypothesis” (Hanslmyr et al., 2012). This model states that lower phase synchronization, not higher, corresponds to a state of higher communication between brain regions. This is due to the supposed more efficient informational transfer capabilities of a desynchronized EEG signal, compared to a synchronized one. In its most recent formulation (Hanslmyr et al., 2016) this theory postulates the existence of two separate phase systems: a cortical system, which communicates through phase desynchronization mainly in the alpha and beta band, and a hippocampal one, mediated by phase synchronization in the theta and gamma band. This hypothesis might explain the aforementioned positive correlations between phase synchronization and dissociative measures, and the negative correlations between synchronization and higher-order functions. Higher synchronization could entail an impoverishment in informational transfer between brain regions, thus producing dissociation effects and loss of higher order functions.

Examining the scalp regions involved in synchronizations related to the state of consciousness, it appears that higher dissociative score correlate with higher alpha, beta and gamma phase synchronization between cortical areas corresponding to the left temporal lobe and right temporo-parietal junction (TPJ), and higher alpha phase synchronization between right temporal lobe and dorsal prefrontal cortex, and right temporal lobe and supplementary motor area (Figure 3, panel A). The activity of the temporal cortices is reported to be strongly involved in dissociative phenomena, both in healthy individuals, in psychiatric patients, and in epileptic patients (Mesulam, 1981). The TPJ play a pivotal role in perceptual integration and self-related processes (Huberle et al., 2012); meditative practices aimed at a “dissolution” of the self can alter the activity of this part of the cortex (Lehmann et al., 2011) and a selective stimulation of this area can produce dissociative symptoms (Orrù et al., 2021). Lesions in the right prefrontal cortex and supplementary motor area might produce asomatognosia (the disappearance of parts of the body from awareness; Arzy et al., 2006), and it is also a known epileptogenic focus in epilepsy patients that report depersonalization (Heydrich et al., 2019). PCI measures of dissociation-like phenomenology, such as Altered State of Consciousness and Altered Experience, also appear to display a convergence with our findings related to dissociative experiences (Figure 3, panel D and E). This altered connectivity between temporal limbic areas and other cortical areas seems to map well with the aforementioned “*Corticolimbic disconnection model”* of dissociation.

In convergence to present findings related to dissociative experiences, we detected opposite correlations between measures relating to phenomenal experiences which are traditionally thought to be reduced in dissociative states. The beta band - volitional control negative correlation topography shown in Figure 4, panel D, linking frontal and central-parietal areas, appears to be in conformity with a modern conceptualization of the beta band as a marker of top-down processes involved in the maintenance of a cortical “*status quo*” (Engel et al., 2010). Furthermore, the positioning of the electrodes displaying significant connections fits well with areas involved in working memory, a cognitive function which is thought to be essential to volition and which has also been found to be mediated by beta band activity (Schmidt et al., 2019). The precuneus, one of the main functional hubs of the default mode network (Marguiles et al., 2009), appears to also be involved. This connection pattern seems to map well with a known functional connection of the precuneus with the dorsal medial and lateral frontal cortex, which has been related to cognitive functions (Zhang & Chiang-shan, 2012). To further point to an involvement of such a frontal-precuneal connection in volition and sleep deprivation, in a recent fMRI study a reduction in functional connectivity between precuneus and middle frontal gyrus has been correlated to attentional decline during acute sleep deprivation (Li et al., 2020).

Beta band analysis also yielded another negative correlation with the experience, much like volition, of a sort of “control” and “order” on thought processes. Attention correlates negatively with phase sync of a dense connection between prefrontal and parietal areas lateralized in the left hemisphere. This topography might suggest an involvement of the fronto-parietal network, which in literature has been tied to sustained attention (Scolari et al., 2015). Interestingly, a previous study has demonstrated that this network displays alterations in its activity in the left hemisphere while shifting from exogenous to endogenous attention. (Meyer et al., 2018). These findings might be ascribable to a reduction in the voluntary component of attention and subsequent increase in dissociative absorption, a known dissociative symptom that can be induced with sleep deprivation (Soffer-Dudek et al., 2018).

## Conclusions

This study empirically demonstrated the use of sleep manipulation to increase dissociative experiences in healthy subjects; the experimental nature of the current investigation could shed further light on the acute nature of dissociative phenomena, and might integrate the well-known “fragmented sleep, fragmented mind” hypothesis (Van Der Kloet et al., 2012), stating the exacerbation of trait dissociation when prolonged disturbed sleep occurs. Pragmatically, these findings might help to better understand the numerous sequelae ascribable to sleep deprivation, generally limited to high order cognitive functions, but poorly characterized in terms of spontaneous cognition. In this regard, dissociative experiences should be interpreted as a regulatory strategy to face the dysfunctional effects of sleep loss, as shown by the association between high EEG bands in frontal regions and dissociative experiences. In this framework, dissociative experiences might be viewed as a compromise to avoid deleterious effects of lack of sleep by generating a unique ineligible state of consciousness in which, according to Tononi’s information integration theory (2008), higher synchronization between brain areas might promote a dissociative processing of information, thus reducing their integration. The results of this dysfunctional state might prevent differentiation between brain modules, resulting in a reduced communication between them. In convergence with Tononi’s theory (2008), when synchronization increases globally, an altered state of consciousness compatible with a dissociative experience could arise from the impoverished information exchange. Contrary to a number of theories of consciousness stating that dissociatives states are prompted by lesser integration and higher segregation within “splitted” modules, our findings indicate that with the “ altered state experiences are positively associated with phase synchronization of high rhythms following which a suboptimal state of consciousness could be driven by an increased synchronization in higher bands, which, in turn, might cause an impoverishment of information exchange between hubs (or modules). Our findings suggest that dissociation could also be prompted not only by the formation of multiple dynamic cores (Lutzenberger et al., 1992) operating with high levels of independence, but also by a low level of segregated and differentiated activity due to high synchronization.

## Material and methods

### Participants

Eighteen healthy volunteers (7 females, age years [28 ± 7]), participated in the study. Subjects eligible for inclusion met the following criteria:

- absence of psychiatric symptoms, verified using the Symptom Checklist-90-Revised (SCL-90-R, Derogatis et al., 1999; Prunas et al., 2011);
- absence of sleep-wakefulness disorders as assessed by the administration and evaluation of Insomnia Severity Index (ISI) (Morin, 1993; Castronovo et al., 2016) and Epworth Sleepiness Scale (ESS; Johns, 1991; Vignatelli et al., 2003) questionnaires
- absence of organic pathologies and of psychotropic addiction verified by an anamnestic questionnaire and by a semi-structured interview conducted by a senior psychiatrist (AG)

Sixteen participants were right-handed, and two were left-handed (Edinburgh handedness inventory, Oldfield, 1971). The study protocol was approved by the Committee on Bioethics of the University of Pisa (Review No. 16/2019 Meeting held on June 28, 2019).

### Experimental protocol

The study consisted of two experimental sessions: a baseline session (i.e., following a whole night of sleep the previous night) and a post sleep deprivation session (i.e., after 36 hours of prolonged wakefulness).

The two sessions were administered in a randomized order at a time distance of one week one from the other, following a restricted randomization procedure (Rosenberger & Lachin, 1988): nine volunteers were submitted first to the baseline and then to the post sleep deprivation session, while the other nine first to the post deprivation and then to the baseline session.

All sessions started at 8 pm and each volunteer was tested individually (i.e. on different days): each participant underwent 10 min of resting state followed by the administration of a battery of psychometric questionnaires to assess state dissociation and phenomenal dimensions of consciousness experienced during the previous time interval. During resting states, participants were lying on a bed in a supine position with eyes closed. Subjects were asked to relax and let their mind wander, avoiding any structured thinking. Resting state’s brain electrical activity was recorded using a high-density EEG system.

Possible confounding factors related to sleep alterations were controlled by monitoring sleep of the nights preceding both baseline and deprivation session: each participant was asked to fill out a sleep diary (Carney et al., 2012; Palagini and Manni, 2016) and night sleep was objectively evaluated using an actigraphic control. Actigraphic control was also used to ensure that volunteers were correctly sleep deprived, since each volunteer spent both the night before the baseline session and the night before the deprivation session at home. The actigraphic detection of any sleep episode during the sleep deprived night implied the exclusion of the volunteer from the study. For this twofold purpose, participants wore an ActiGraph wGT3X-BT (ActiGraph, Pensacola, FL, USA) placed on their non-dominant wrist. Actigraphic data were analyzed and visually inspected using Actilife software (version 6.11.9).

### Psychometric questionnaires

#### Psychometric assessment of the eligibility criteria

To assess the eligibility of volunteers, we administered the following questionnaires for psychological symptoms and sleep disorders:

The Symptom Checklist-90-Revised (SCL-90-R) scale comprising 90 items (Derogatis et al., 1999). SCL-90-R is a self-administered questionnaire evaluating the patterns of psychological symptoms. The results are shown as nine domain scores of primary dimensions including somatization, obsessive-compulsive, depression, interpersonal sensitivity, anxiety, hostility, phobic anxiety, paranoia, and psychotic symptoms and three global indices, namely the global severity index, the positive symptoms total, and the positive symptoms distress index. Each question is scored based on a 5-point Likert scale from 0 (none) to 4 (extreme). The Italian adaptation of the questionnaire was used (Sarno et al., 2011). All participants scored below the cut-off in both primary dimensions and global indices

The Epworth Sleepiness Scale (ESS) (Johns, 1991) a self-administered questionnaire to measure a subject’s general level of daytime sleepiness. The ESS comprises eight items evaluating the subject’s likelihood of sleepiness or falling asleep in a particular situation in daily life situations, and the total scores measures the subject’s average sleep propensity across those different situations in daily life. Each question is scored based on a 4-point scale for each of the eight questions. The Italian adaptation of the questionnaire was used (Vignatelli et al., 2003). All participants scored below the cut-off, thus evidencing the absence of sleep propensity.

The Insomnia Severity Index (ISI) is a seven-item self-administered questionnaire assessing the nature, severity, and impact of insomnia within the last month (Morin, 1993). The dimensions evaluated comprises the severity of sleep onset, sleep maintenance, and early morning awakening problems, sleep dissatisfaction, interference of sleep difficulties with daytime functioning, noticeability of sleep problems by others, and distress caused by the sleep difficulties. Each question is scored based on a 5-point Likert scale (0 = no problem; 4 = very severe problem), yielding a total score ranging from 0 to 28. The total score is interpreted as follows: absence of insomnia (0–7); sub-threshold insomnia (8–14); moderate insomnia (15–21); and severe insomnia (22–28). Three versions are available—patient, clinician, and significant others—but the present paper focuses on the patient version only.

The Italian adaptation of the questionnaire was used (Castronovo et al., 2016). All participants scored below the cut-off.

### Descriptive statistics of inclusion questionnaires are reported in the *supplementary materials*

#### Clinician Administered Dissociative State Scale

The Clinician Administered Dissociative State Scale (CADDS), developed by Bremner et al. (1998), is a psychometric questionnaire quantifying present-state dissociation. It consists of 19 self-report items, (scored as 0 = not at all; 1 = slightly; 2 = moderately; 3 = considerably; 4 = extremely), which yield three subjective subscale scores (amnesia, depersonalization, and derealization) and of 8 clinician rated items that result in one observer rated score. The CADSS-total score can range from 0 to 76. Subscales of CADSS provide measures of the following dissociative phenomena: depersonalization (items 3–7), derealization (items 1, 2, 8–13, 16–19), and dissociative amnesia (items 14,15). The Italian adaptation, avalaible at https://eprovide.mapi-trust.org/instruments/clinician-administered-dissociative-states-scale, has been administered.

#### Phenomenology of Consciousness Inventory

The Phenomenology of Consciousness Inventory (PCI, Pekala, 1991) is a retrospective, phenomenologically-based self-report instrument quantifying subjective experiences by assessing 12 major dimensions of consciousness: altered experience; positive affect, negative affect, attention, imagery, self-awareness, state of awareness, internal dialogue, rationality, volitional control, memory and arousal. In the present study, the Italian version of PCI, available at www.quantifyingconsciousness.com, has been administered (Pekala et al., 2017).

### EEG recording and preprocessing

Electroencephalographic recordings were carried using a Net Amps 300 system (GES300) and a 128 electrode HydroGel Geodesic Sensor Net (ElectricalGeodesic Inc., Eugene, OR, USA) plus two ECG electrodes, one placed over the right clavicle and the second on the lower left rib cage, ECG derivation was used as reference for cardiac artifacts.

Signals were acquired referenced to the vertex at a 500 Hz sampling rate, keeping electrodes impedances below 50kΩ, using Net Station software (version 4.4.2). EEG signals were preprocessed using EEGLAB (Delorme and Makeig 2004), a Matlab toolbox (MathWorks, Natick, MA, USA) for processing continuous and event-related electrophysiological data. Signals were down-sampled from 500 to 125 Hz and band-pass filtered between 1 Hz and 45 Hz (two-way least-squares FIR filtering).

Channels located on the forehead, cheeks and nape, which mainly contribute to movement-related noise, were discarded (Chennu et al., 2014), thus retaining 99 channels out of 128. Retained signals were independently visually inspected by two EEG experts (VC and EC) to reject segments affected by non-stereotyped artifacts such as subject’s movements (i.e., swallows, sudden head or body movements), electrode cables movements or other external sources of interference. Artifacts caused by eye movements, eye blinks, muscle tension, and heart pulse were kept for further preprocessing: due to their stereotyped nature, these artifacts can be reliably identified and separated into only a few independent components using the Independent Component Analysis (ICA; Makeig et al., 1996) and then discarded to obtain putatively artifact-free EEG signals. The selection of artifactual components was based on a visual examination of each component time course and power spectrum, as well as on the analytic tools available from the ICLabel toolbox (Pion-Tonachini et al., 2019) to support visual inspection. The ICLabel component rejection threshold intervals were set between 0-0.15 for “Brain” classifications, and between 0.8-1 for all others (Muscle, Eye, Heart, Line Noise, Channel Noise, Other). The ECG signal was used as a reference for cardiac artifact identification. For the following analysis, EEG signals were finally re-referenced to the mastoids’ average signals (Massimini et al., 2004).

### EEG band spectral content

Power analysis associated with EEG oscillations is the basic approach to identifying cortical rhythms generated by the activity of specific functional circuits, and to assess the degree of cortical recruitment (Schomer and Da Silva, 2012). Thus, a power spectrum analysis was conducted for each subject and session (baseline and post deprivation) The mean power spectrum density (PSD), as a function of frequency was estimated for each EEG channel by applying a sliding (50% overlap between consecutive epochs). Hamming-windowed Fast Fourier Transform (FFT) on 4 sec epochs and averaging between them. Band-wise PSD at each channel was then evaluated for five bands of interest: delta (1–4 Hz), theta (4–8 Hz), alpha (8–12 Hz), beta (12–30 Hz) and gamma (30–45 Hz), by averaging among the bins pertaining to the band and log-transforming the mean value (dB).

### EEG phase synchronization

Phase synchronization, widely regarded as an EEG measure of information exchange between neuronal populations, is often calculated examining the time course of phase differences between signals at different electrodes couples.For each subject, condition (baseline and post deprivation) and band of interest, connectivity between each couple of electrodes was estimated using the debiased Weighted PLI (dwPLI; Vinck et al., 2011). DwPLI belongs to the family of phase synchronization estimators that minimize the effects of volume conduction, common sources, and noise (Stam et al., 2007; Lachaux et al., 1999).

DwPLI at frequency bins of 0.25Hz (range 1-45 Hz) was estimated on 4 sec epochs (50% overlap between consecutive epochs). For each epoch and couple of electrodes, the dwPLI of each band of interest was obtained by averaging over the dwPLI values of its frequency bins. The average dwPLI for each subject, session, band and couple of electrodes was finally estimated by averaging over the epochs pertaining to the session.

### Statistical analysis

#### Psychometric tests

Putative between-conditions (post-deprivation versus baseline) differences were evaluated for each psychometric measure (CADSS total score and its three subdimensions scores as well as PCI dimensions scores), using the paired Wilcoxon Signed rank test (significance threshold was set at p ≤ 0.05). The magnitude of between-conditions changes was assessed by estimating the effect size for non-parametric tests (Fritz et al., 2012).

Furthermore, after singling out eventual significant changes in psychometric measures, we explored their EEG correlates.

The Spearman’s rank correlation test was used to estimate associations between changes from baseline to post deprivation of each psychometric measure responsive to sleep deprivation and those of EEG features. Let us assume without loss of generality to consider a psychometric variable *x* and an EEG feature *y* estimated at one electrode (PSD) or couple of electrodes (dWPLI); the Spearman correlation test is conducted between *Δx = x*_*post deprivation*_ *- x*_*baseline*_ *and Δy = y*_*post deprivation*_ *- y*_*baseline*_.

Correlation significance at each electrode (PSD) or couple of electrodes (dWPLI) was then assessed using a single threshold permutation test for the maximum r-statistic with 10000 randomizations (Statistical nonParametric Mapping, SnPm, Nichols and Holmes, 2002).

SnPM approach was chosen to counteract the multiple testing issues which arise when considering simultaneous testing on dense electrode arrays (99 electrodes/tests for band-wise PSDs and 4851 couple of electrodes/tests for band-wise dWPLI), as (1) it does not require any assumption on data normality, and (2) is a simple yet robust approach to control for type I statistical errors (i.e., rejection of a true null-hypothesis). The significance threshold was set at p ≤ 0.05 after multiple testing corrections.

Finally, between-conditions comparisons (using paired t-tests), were conducted for those EEG features showing significant associations with psychometric measures. For each set of comparisons (either band-wise PSDs or band-wise dwPLI), t-value significance at each electrode was estimated using a single threshold permutation test for the maximum t-statistic with 10000 randomizations (Nichols and Holmes, 2002). Also in this case, the significance threshold was set at p ≤ 0.05 after correction for multiple testing.

All statistical analyses were conducted using tailored codes written in Matlab (MathWorks, Natick, MA, USA) and Statistical Package for the Social Sciences (SPSS statistics, IBM SPSS, IBM Corporation).

## Notes

### Competing Interest Statement

The authors have declared no competing interest.

