## Supplemental tables for "Sleep Deprivation Induces Acute Dissociation via Altered EEG Rhythms Expression and Connectivity"

**Table S1: Inclusion questionnaires**

| Questionnaire | Dimension | Mean ± SD |
|---|---|---|
| Insomnia Severity Index | TOTAL SCORE | 4,3 (± 2,12) |
| Epworth Sleepiness Scale | TOTAL SCORE | 6,5 (±4,23) |
| Symptoms Checklist 90-Revised | SOM | 44,35 (± 5,12) |
| | O-C | 40,76 (± 4,07) |
| | I-S | 43,06 (± 5,71) |
| | DEP | 42,41 (± 3,48) |
| | ANX | 44,29 (± 3,71) |
| | HOS | 43,59 (± 5,79) |
| | PHOB | 44,76 (±1,99) |
| | PAR | 42,47 (± 6,17) |
| | PSY | 43, 76 (±4,88) |
| | GSI | 42,12 (± 4,07) |
| | PST | 41,06 (± 6,48) |
| | PSDI | 42,65 (± 5,48) |

**Table S1:** Descriptive statistics (mean ± SD) for each inclusion self-administered questionnaire. For ISI and ESS, the total score is well below the critical cut-off (ESS< 15; ISI < 15); For SCL 90-R, both primary dimensions and global indices are well below the critical cut-off <50.
ANX, anxiety; DEP, Depression; ESS, Epworth Sleepiness Scale; GSI, Global Severity Index; HOS, Hostility; I-S, Interpersonal Sensitivity; ISI, Insomnia Severity Index; O-C, Obsessive-Compulsivity; PAR, paranoid ideation; PSY, Psicoticism; PHOB, Phobic anxiety; PS, Positive Symptom Total. PSDI, Positive Symptoms Distress; SCL-90-R, Symptoms Checklist 90- Revised; Som, Somatization;

**Table S2: Phenomenological Consciousness Questionnaire (Including Minor Dimensions)**

| Name | Sum Of Ranks | z Score | P value |
|---|---|---|---|
| **Positive Affect** | 99 | 1,609711009 | 0.107 |
| *Joy* | 29,5 | -0,7507477407 | 0.512 |
| *Sexual Arousal* | 24 | 1,705605731 | 0.109 |
| *Love* | 85,5 | 2,085019278 | 0.035* |
| **Negative Affect** | 49 | 0,2450490147 | 0.827 |
| *Anger* | 21,5 | -0,1187827742 | 0.945 |
| *Sadness* | 35 | 1,491374966 | 0.160 |
| *Fear* | 8,5 | 0,2709141846 | 0.875 |
| **Body Image** | 28 | -0,8642416215 | 0.423 |
| **Altered State Of Awareness** | 26 | -2,394602324 | 0.017* |
| **Altered Experience** | 26,5 | -2,14663149 | 0.032* |
| *Altered Time Sense* | 28 | -1,611980992 | 0.107 |

| | | | |
|---|---|---|---|
| *Altered Perception* | 14 | -2,425272035 | 0.012* |
| *Altered Meaning* | 57,5 | -0,5466081666 | 0.585 |
| **Imagery** | 97 | 0,9735724153 | 0.330 |
| *Imagery Amount* | 57,5 | 0,4756514942 | 0.669 |
| *Imagery Vividness* | 84 | 1,374517695 | 0.184 |
| **Attention** | 97 | 1,9373298 | 0.052 |
| *Direction Of Attention* | 84,5 | 1,396867687 | 0.172 |
| *Absorption* | 84,5 | 2,094051328 | 0.036* |
| **Self-Awareness** | 153 | 3,627466888 | 0.000*** |
| **Internal Dialogue** | 122 | 2,167588456 | 0.030* |
| **Rationality** | 133,5 | 2,703198368 | 0.007** |
| **Volition** | 142 | 3,110682595 | 0.002** |
| **Memory** | 147,5 | 3,366195602 | 0.001*** |
| **Arousal** | 71 | 1,16596896 | 0.258 |
